# A sparse set of spikes corresponding to reliable correlations is highly informative of visual stimulus on single trials

**DOI:** 10.1101/2022.01.24.477564

**Authors:** Maayan Levy, Jeff K. Guo, Jason N. MacLean

## Abstract

Spike trains in cortical neuronal populations vary in number and timing trial-to-trial, rendering a viable single trial coding scheme for sensory information elusive. Correlations between pairs of neocortical neurons can be segmented into either sensory or noise according to their stimulus specificity. Here we show that pairs of spikes, corresponding to reliable sensory correlations in imaged populations in layer 2/3 of mouse visual cortex are particularly informative of visual stimuli. This set of spikes is sparse and exhibits comparable levels of trial-to-trial variance relative to the full spike train. Despite this, correspondence of pairs of spikes to a specific set of sensory correlations identifies spikes that carry more information per spike about the visual stimulus than the full set or any other matched set of spikes. Moreover, this sparse subset is more accurately decoded, regardless of the decoding algorithm employed. Our findings suggest that consistent pairwise correlations between neurons, rather than first-order statistical features of spike trains, may be an organizational principle of a single trial sensory coding scheme.

---

In order to meet the demands of survival, organisms must be able to process and act upon intrinsic (i.e. attention, hunger) and extrinsic (i.e. sensory stimuli) variables simultaneously. Indeed, both intrinsic^1,2^ and extrinsic^3^ variables contribute to action potential generation or spikes in neocortical neurons. Spikes and the intervals between them can be considered the symbols of a coding scheme. At the single neuron level, the scheme has been postulated to depend on the number of spikes emitted, i.e. rate code^4,5^ and/or the specific sequence of spikes and silences, i.e. a temporal code^6–8^. In general, when evaluating a coding scheme, the focus has been on trial averaged spiking activity. However, spike counts and sequences of spikes are highly variable trail-to-trial^9–13^, rendering an individual neuron-based coding scheme that is viable for single trials elusive. When considering a population of neurons there are numerous combinatorial possibilities involving both rate and timing, including but not exclusive to relative spike times^14,15^ and covariance of spike counts^16^. Despite these advances, a population level, single trial coding scheme remains unclear. The challenge of delineating a single trial coding scheme is further exacerbated by the difficulty in disambiguating the relative contribution of intrinsic and extrinsic variables to spiking.

Spike trains within a population of neocortical neurons exhibit pairwise and higher order correlations (for reviews see^17–19^). Pairwise correlations, like spikes, arise from variables both external and internal to the population of neurons recorded. Unlike spikes, studies have successfully segmented correlations demonstrating that correlations can be stimulus dependent^20–22^, can reflect local integration of synaptic inputs^23,24^, and can arise from global or broadcast signals within neocortex^25,26^. Pairwise correlations have been shown to contain sufficient information about neuronal dynamics to accurately predict single trial activity of a given neuron from its correlated counterparts^27–29^, termed peer-prediction^30^. Moreover, pairwise correlations have also been employed to construct accurate single trial decoders^20,28^. Both lines of evidence indicate that pairwise correlations carry information about sensory stimuli above and beyond that carried by spikes alone^27,31^. While any one spike is clearly the consequence of the combination numerous inputs, and by extension variables, the fact that correlations can be segmented according to variable presented the possibility that pairs of spikes corresponding to pairwise correlations can similarly be segmented, presumably revealing the main contributing variable. To do so, we sort spikes on individual trials, according to whether they correspond to stimulus or non-stimulus specific pairwise correlations, and then use information theoretics and decoders to test our hypothesis.

We find that a sparse set of stimulus-specific spikes, embedded within the full and denser single trial spike train, carries more information per spike about the visual stimulus than the full set or any other matched set of spikes. Moreover, this sparse subset is more accurately decoded, regardless of decoding algorithm employed. Our results suggest that consistent pairwise correlations between neurons across trials, rather than specific first-order features of spike trains, may be an organizational principle of a single trial sensory coding scheme. Such a scheme allows for spike trains within individual neurons to potentially represent both sensory, global and local variables simultaneously and that the associated pairwise correlations, and potential diversity of post-synaptic targets, allow for each variable to be distinguishable from each other downstream.

## Results

We imaged hundreds of excitatory neurons in layer 2/3 of mouse primary visual cortex (V1) in response to 12 directions of drifting gratings^32^. We summarized the spiking dynamics of these populations of neurons as functional networks (FNs), with neurons and the correlations between them as nodes and edges, respectively. We distinguished globally driven, from stimulus-specific, correlations by computing a functional network for each of the 12 directions of drifting gratings using a mutual information measure (conMI^33^; Fig. 1a) between each pair of neurons in all trials of each grating direction separately. conMI is the mutual information between neuron *i* at time *t* and neuron *j* at time *t* and *t+1*, with time dictated by imaging frame.

**Figure 1.**
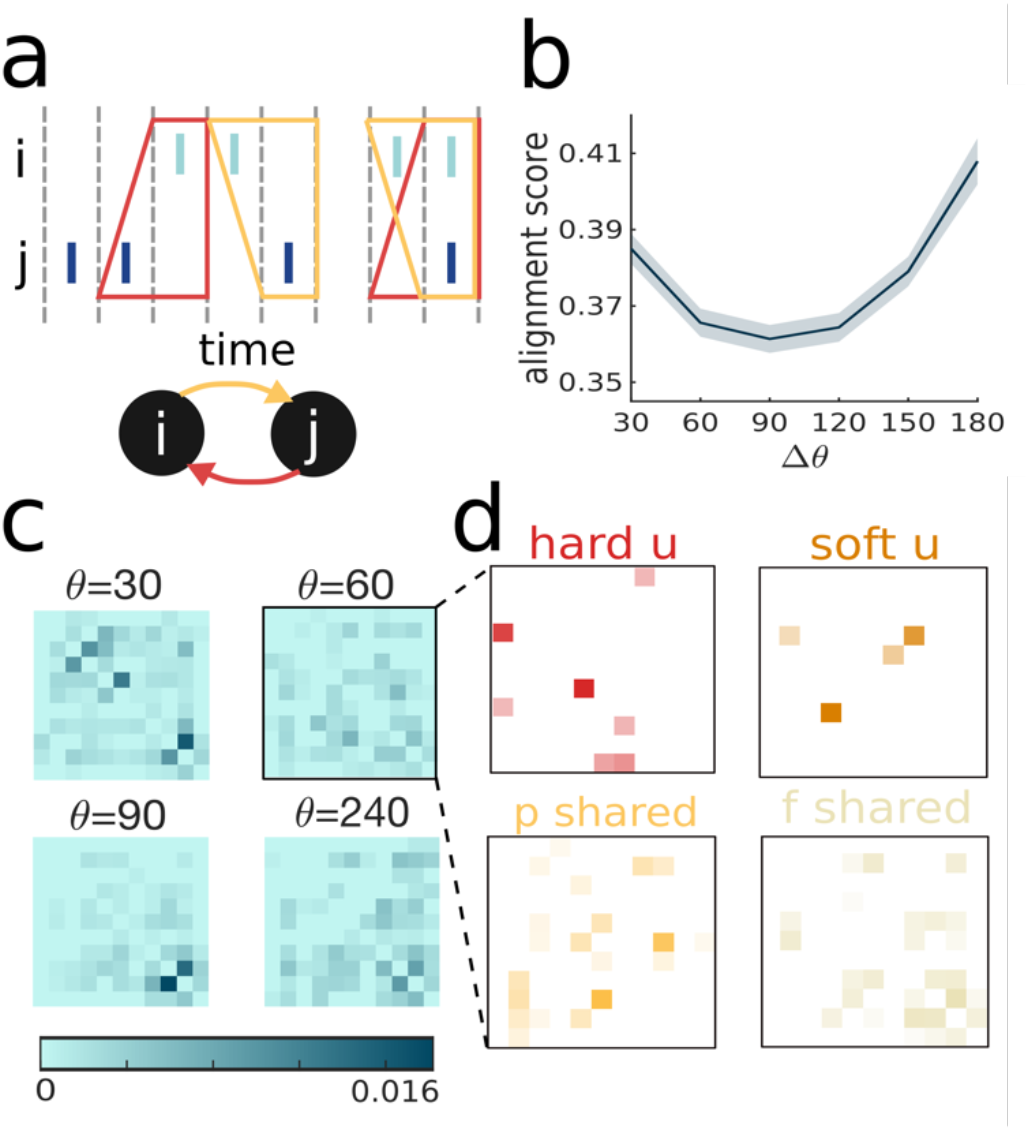
functional networks (FNs) can be divided to 4 sub-FNs based on edge statistics. **(a)** Illustration of edge inference by confluent mutual information (conMI). The edge from neuron *i* to neuron *j* is a statistical dependency between *i* spiking at time *t-1* or *t*, and *j* spiking at time *t*. Hence conMI is not necessarily symmetric. **(b)** FN similarity reflects stimulus similarity: alignment score varies in [0,1] with larger numbers indicating increased similarity between two networks, and is computed as in ref 20. Δ*θ* stands for stimulus similarity in degrees, with 30 degrees being an adjacent direction and 180 degrees being the direction with the same orientation. Line and shading represent mean and the standard error across datasets, respectively. **(c)** FNs for 4 directions: the direction of interest (60 degrees) and its three fellow directions (30,90 and 240 degrees). 10 neurons are illustrated here for visualization purposes. Colors represent edge weight. Note that some edges are similar across directions whereas others appear only in one FN. **(d)** Illustration of edges allocation into 4 sub-FNs: the FN for 60 degrees from B was split into 4 non-overlapping sub-FNs, according to which edges are unique and strong in the direction of interest (60 degrees) as compared to the FNs for fellow directions. Colors for sub-FNs are consistent throughout this manuscript.

Next, we systematically compared edges in the twelve FNs. We observed that some edges in the FNs are overlapping across stimuli, whereas other edges are present only in a subset or a single FN corresponding to a single direction of drifting gratings (Fig. 1b)(11.53±1.62% unique edges, 12.96±2.66% edges shared between two directions, 75.51±3.98% edges that are present in more than 2 FNs). Furthermore, the similarity of FNs correlated with stimulus similarity^20^ (Fig. 1c), with increased similarity for directions that are adjacent (e.g. 30 and 90 are adjacent to 60) and directions that have the same bar orientation (e.g. 60 and 240). This finding suggests that these three (fellow) directions are potentially more difficult for downstream circuits to disambiguate. We allocated edges in each FN (termed the overall FN throughout the manuscript) into four sub-FNs depending on the extent to which an edge was shared or unique to one FN corresponding to one direction of drifting gratings. Specifically, edges were sorted into: *hard u(nique)* sub-FN which contains edges that only exist in an FN for a single direction, *soft u(nique)* edges may exist in FNs for fellow directions, but they are stronger, reflecting more reliable statistical dependency, in an FN for one direction. *P(artial) shared* which consists of edges that are common to a direction and to some, but not all fellow directions, and *f(ully) shared* edges that are found in all the FN regardless of direction. This sorting procedure was done for each direction of drifting grating and resulted in 4 non-overlapping sub-FNs (Fig. 1d).

Across datasets and directions, 17.41±5.06% edges were classified as *hard u*, 6.89±2.49% edges as *soft u*, and 53.46±4.31%, 22.22±9.02% edges as *p shared* and *f shared*, respectively, rendering *soft u* the sparsest sub-FN and *p shared* the densest sub-FN (Fig. 2a). In other words, we find that a minority of correlations are specific to the sensory stimulus. As expected, given our edge segmentation procedure, we found that *soft u* contains a significant subset of the strongest edge weights (AVONA, p<0.001, Fig. 2b), meaning more of the edges of this sub-FN reflected reliable statistical dependencies as compared to the other three sub-FNs. Notably, the sub-FNs included most of the recorded neurons (*hard u*: 99.39±0.99%, *soft u*: 95.15±6.06%, *p shared*: 99.15±1.14% and *f shared*: 91.80±4.17%) demonstrating that the vast majority of all neurons had at minimum one edge in every sub-FN, ruling out a scheme in which a neuron codes for **either** sensory **or** global variables.

**Figure 2.**
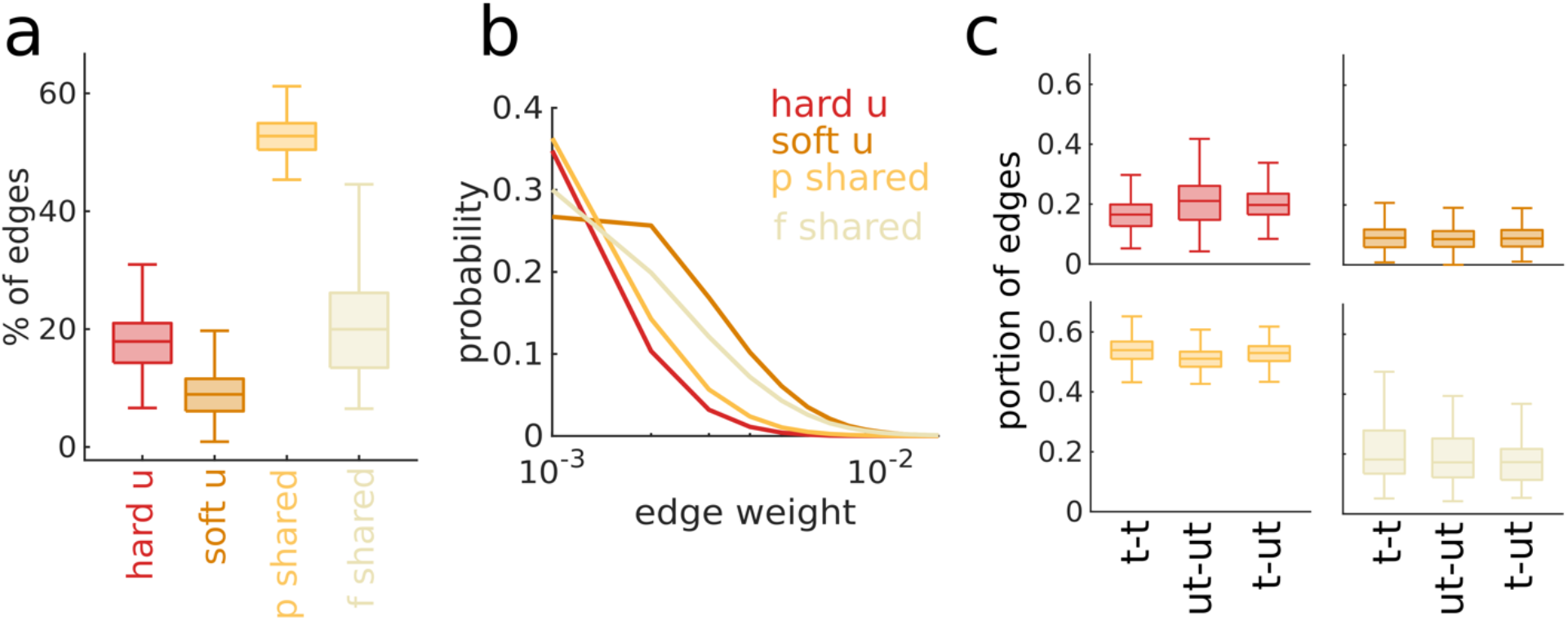
topological properties of sub-FNs. Data in this figure are shown across datasets and stimuli unless stated otherwise. **(a)** Density of the four sub-FNs as the portions of edges from the overall allocated to each of them. Boxplots represent interquartile ranges and midlines mark the median. **(b)** Probability distributions for the edge weights included in each of the four sub-FNs. **(c)** Portion of edges between tuned-tuned (t-t), untuned-untuned (ut-ut) and mixed (t-ut) neurons for the four sub-FN out of the total number of edges in the overall FN. Boxplots and midlines stand for the interquartile range and the median, respectively.

Since functionally similar neurons have been reported to be more likely synaptically connected as well as correlated^32,34,35^, we next evaluated the extent to which the edges within each sub-FN may simply be due to similar tuning of individual neurons within the FN. We counted how many edges in each sub-FN are between pairs of tuned (t-t), untuned (ut-ut), and mixed-functionality (t-ut, ut-t) neurons. We found that the prevalence of each edge type was proportional to the density of the sub-FNs (Fig. 2c), indicating that correlations between tuned neurons are not over or under represented in sub-FNs.

### Spikes that correspond to sub-FNs are sparse and variable

From the perspective of a theoretical downstream neuron, correlated synaptic inputs from multiple pre-synaptic neurons are far more likely to result in cooperative integration and result in post-synaptic neuronal spiking and in turn effective information transfer^23,36^. Adopting the perspective of a downstream neuron, we segmented spikes according to correspondence to an edge in a sub-FN. In other words, if neuron *i* spiked at time *t* and neuron *j* spiked at time *t+1*, we kept those spikes if there was a non-zero edge in the *ij*-th index in the FN, and discarded spikes that did not correspond to an edge (Methods; Fig. 3a). This procedure yielded 4 sets of spikes that corresponded to edges in each sub-FN for each trial. Note that no new spikes are inserted. Rather, we picked subsets of the original spike trains. We call these subsets of spikes sub-FN consistent spikes.

**Figure 3.**
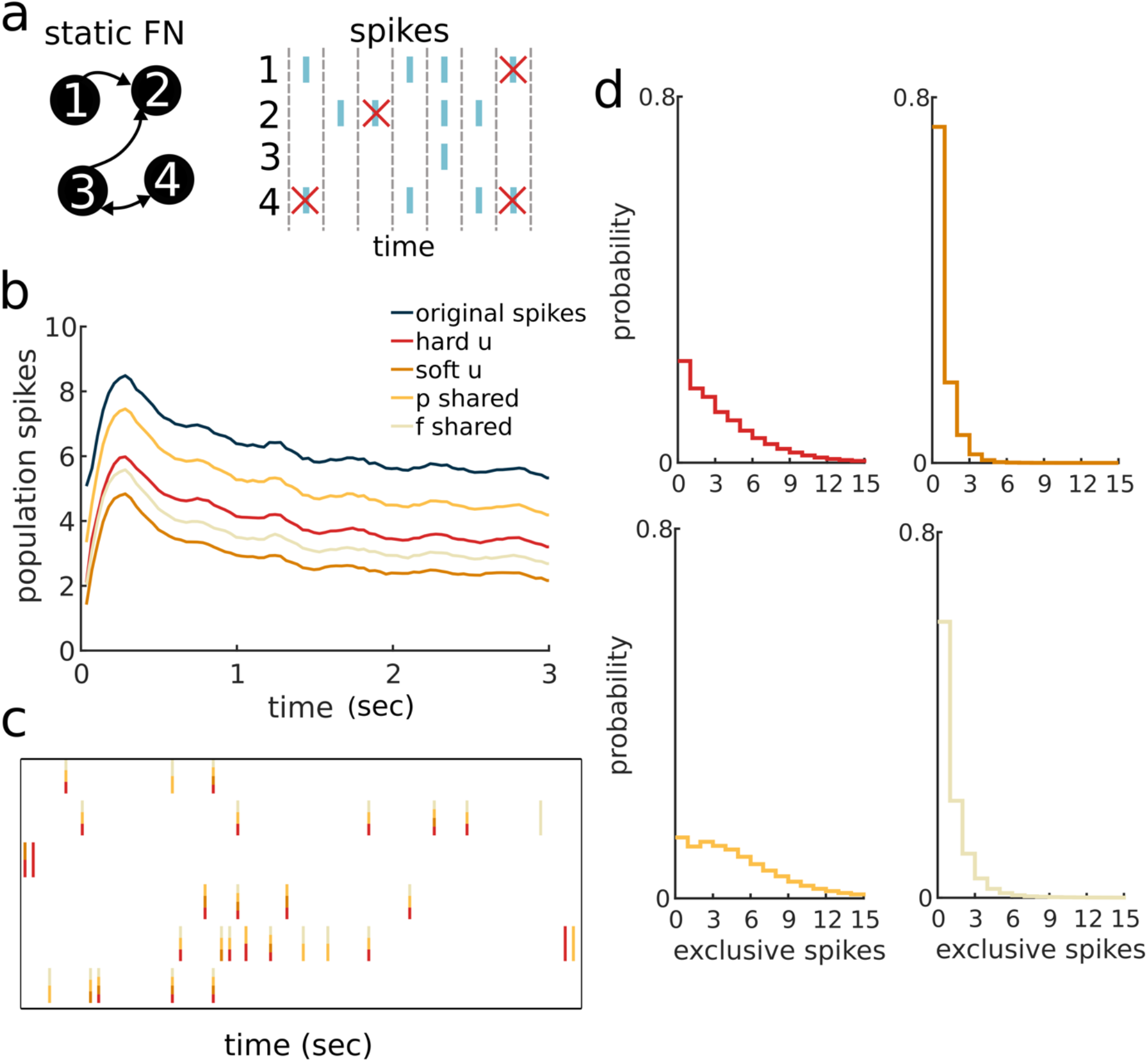
sub-FN consistent spikes are ultra-sparse and non-exclusive. **(a)** Illustration of a temporal graph (TG) approach: a static FN which is inferred from many trials is intersected with spikes on a single-trial time-point by time-point basis. An edge from neuron 1 to neuron 2 is expressed in the first two time-frames with neuron 1 spiking followed by a spike in neuron 2. Any static FN can be used in this procedure, and we have used both the overall FN and the sub-FNs. Spikes that are not an expression of any edges are marked with a red X and later discarded in the spike sparsification process. **(b)** Population spikes over time in the trial for the original spikes and the four sets of sub-FN consistent spikes. Lines represent means across datasets. **(c)** Raster plot of 6 randomly sampled neurons from one randomly sampled dataset. Spikes are colored according to the sub-FN they are consistent with. For example, a spike colored in red and yellow is consistent with both hard u and p shared. **(d)** Probability distributions of spikes that are consistent only with one sub-FN (i.e. exclusive) in one time point. Data across datasets and trials.

Sub-FN consistent spikes are sparser than the original population singe trial spikes, with *soft u* corresponding to the fewest action potentials (Fig. 3b). Notably, in this framework, a spike can correspond to an edge in more than one sub-FN; for example, an *ij* edge may exist in *hard u*, while an *ik* edge is nonzero in *p shared*. In this case, a spike at time *t* will be retained for both of these sub-FNs if neurons *j* and *k* spiked at time *t+1*, respectively. Therefore, while edges are exclusive to one of the four sub-FNs, the corresponding spikes are not. We found varying degrees of overlap between sub-FN consistent spikes for different sub-FNs (Fig. 3c-d).

Sub-FN consistent spikes within individual neurons were no less variable trial-to-trial, relative to the full spike trains, as measured by rate or temporal precision (Fig. 4a and 4c, respectively). Similarly, we found that network wide sub-FN consistent spikes were not more reliable than the original spikes as a population vector as measured with the L2 norm (Fig. 4b).

**Figure 4.**
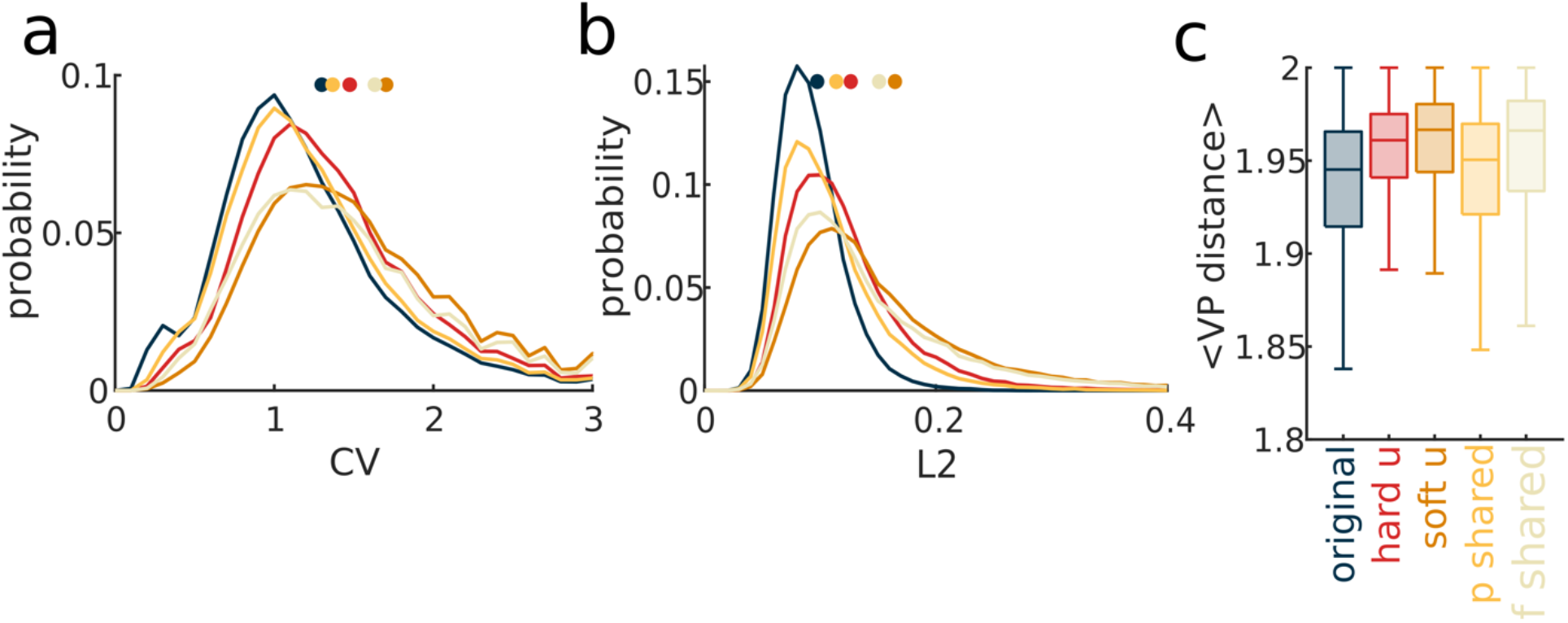
trial-to-trial variability of sub-FN consistent spikes. Data in this figure are shown across datasets and stimuli. **(a)** Coefficient of variation probability distributions for the firing rate of single neurons in the original spikes and in each of the sets of sub-FN consistent spikes. Dots on the top indicate the means. Colors as in (c). **(b)** L2 (Euclidean) norm (see Methods) probability distributions for population vectors in the original spikes and in each of the sets of sub-FN consistent spikes. Dots on the top indicate the means. Colors as in (c). **(c)** Temporal precision as measured by the VP distance. We normalized the metric by the number of spikes (Methods). <> denotes the mean over pairs of trials of the same grating direction. Boxes span the interquartile range and horizontal lines indicate the medians.

### *Soft u* consistent spikes are more informative of drifting grading direction

Information theory has been used to great effect in neuroscience to examine coding in neural systems^37,38^ and offers a direct measure of the information present in neural responses about a stimulus. Moreover, previous work in cat LGN has shown that pairs of spikes between neurons, like those considered here, carry more information about a visual stimulus as compared with the same numbers of unpaired spikes^39^. To quantify information in the full spike trains and in sub-FN consistent spikes we calculated the mutual information between spikes and stimulus (Methods).

We normalized the raw information measure by the average number of spikes to obtain a metric of bits per spike regardless of sparsity. We examined the information in a small group of neurons: we picked 5 neurons that had the largest average firing rates in the full set of spikes across stimuli and trials, and held the identities and order of these 5 neurons fixed. We selected the high firing rate neurons to ensure enough spikes to compute information since each subset of spikes was very sparse.

Examining the activity in bins of 10 frames, we found that *soft u* consistent spikes were significantly more informative about the direction of drifting gratings (1.07±0.56 bits/spike), as compared to the original spikes (0.13±0.15 bits/spike) and other subsets of spikes (0.48-0.67±0.26-0.5 bits/spike) (Fig. 5a). To quantify the information on faster timescales, we concatenated every 3 imaging frames preserving the spatio-temporal pattern of spikes. Overall, we observed more information per spike in these spatio-temporal patterns as compared to binned frames and again *soft u* consistent spikes contained more information per spike (1.75±0.5 bits/spike) as compared to all of the action potentials (0.99±0.13 bits/spike) and to the other sets of sub-FNs consistent spikes (1.27-1.38±0.18-0.49 bits/spike) (Fig. 5b). Notably, quantifying the information in omitted spikes, that is, spikes that were *not* consistent with sub-FN edges resulted in the opposite trend with *p shared* leading in information per spike (Fig. 5d) consistent with the little overlap between *soft u* and *p shared* consistent spikes.

**Figure 5.**
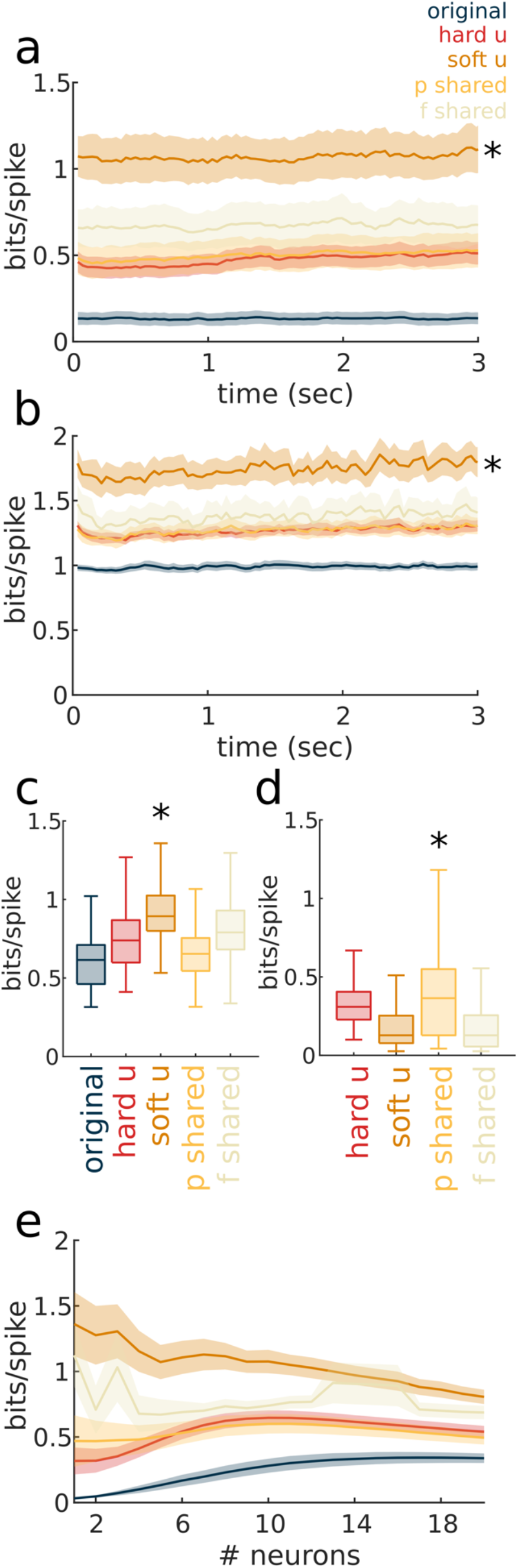
information quantity in sets of sub-FN consistent spikes. All panels show data across datasets. **(a)** Bits per spike in sub-FN consistent spikes (different sub-FNs in their respective colors) and original spikes across time. Here we quantified the information in “binary words” of size 5, using the 5 neurons with the largest firing rate and binning over 10 frames. Line and shading represent the mean and standard error, respectively. *p<0.05, ANOVA test for every timepoint. *Soft u* is significantly different from all other line except f shared. **(b)** Bits per spike in sub-FN consistent spikes (different sub-FNs in their respective colors) and original spikes across time. Here we quantified the information in spatio-temporal patterns where every 3 consecutive frames were vectorized, which preserves the structure of spiking in the 5 neurons we analyzed. *p<0.05, ANOVA test for every timepoint. *Soft u* is significantly different from all lines as all times, except *f shared* (p<0.05 only for 32/100 time points). **(c)** Bits per spike in sub-FN consistent spikes (different sub-FNs in their respective colors) and original spikes in a group of 5 neurons with the firing rates closest to the population mean. Boxplots and midlines denote interquartile range and medians across all time points. *p<0.01 **(d)** The same as (a) and (b) but for spikes that are not consistent with the sub-FN. For example, for *hard u* we used spike that are discarded when intersecting *hard u* sub-FN with the spikes. Boxplots and midlines represent interquartile range and medians across all time points, respectively. *p<0.05, except for *soft u-f shared* comparison. **(e)** Information per spike as a function of the number of neurons included in the analysis. Lines and shading are for the mean and standard error, respectively.

We next asked whether neurons which exhibited firing rates comparable to the population mean firing rate, rather than neurons with the highest firing rates showed similar results. *Soft u* consistent spikes again carried more information about the direction of drifting grating in these groups of neurons (Fig. 5c). However, we found that as larger and larger groups of neurons are included in the calculation that *soft u* consistent spikes offer little extra information beyond that contained in small groups (Fig. 5e) indicating greater redundancy in the *soft u* consistent spikes. In contrast, other subsets of spikes followed uniform profiles, where larger pools sum up linearly, and the full, original spikes are more informative as more neurons are included (Fig. 5e).

### *Soft u* consistent spikes are decodable at high accuracy

The elevated information quantity in *soft u* consistent spikes suggests that these spikes may be more decodable. To evaluate whether this is the case we used a feedforward network multiclass decoder trained with conjugate gradient on 90% of the data yielded significantly higher performance on the test set for *soft u* consistent spikes (52.13±14.60%) as compared to all action potentials and all of the other subsets of spikes (34.12±11.8% for the original spikes. 44.37±12.80%, 40.64±11.20% and 42.95±13.06% for *hard u, p shared* and *f shared*, respectively, Fig. 6a).

**Figure 6.**
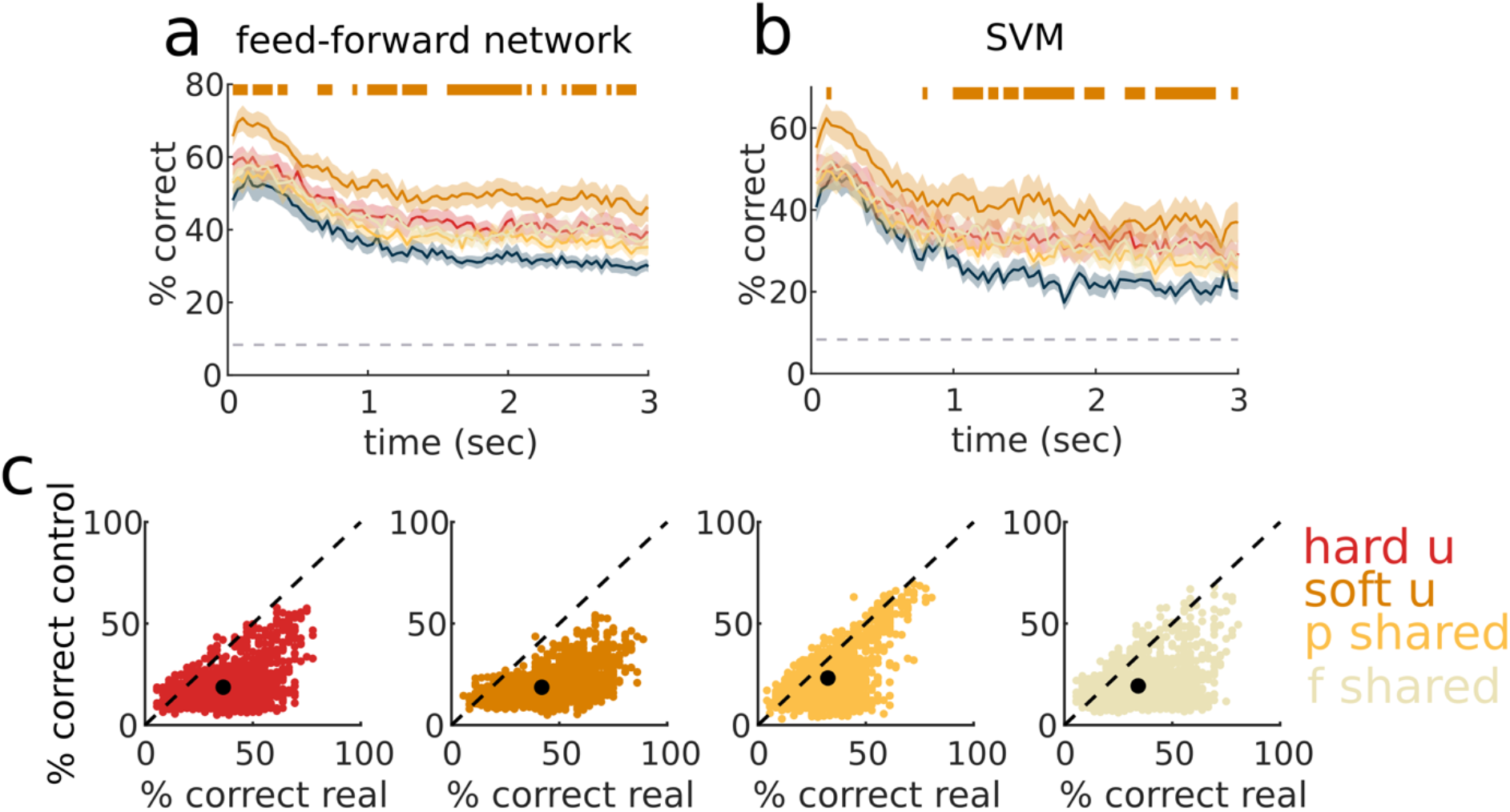
decoding the direction of drifting gratings from sub-FN consistent spikes. **(a)** Performance as percent correct from the test set trials of the original spikes and the sparsified sub-FN consistent spikes in the Feedforward neural network decoder (Methods). Dashed line on the bottom denotes chance level decoding (1/12). The original spikes are in blue. Orange bars on top indicate the time points at which *soft u* is significantly better (ANOVA, p<0.05). Lines and shading are for the means and standard errors across 16 datasets. **(b)** Performance of a second supervised decoder, namely a Support vector machine (SVM). Colors and markings are the same as in (a). **(c)** Real decoding performance (abscissa) of the four sets of sub-FN consistent spikes against the decoding performance of control spikes (ordinate) in the SVM framework. Control spikes were generated by permuting the sub-FN before intersection with the original spikes and thus preserve the sparsity and the same count of pairs of consecutive spikes but not their specificity (see Methods). Dashed line is the identity line. Each data point is one decoder, binned at 10 frames. Black dots represent cloud means.

To ensure that this decoding performance did not depend on decoder architecture we confirmed these findings with a support vector machine (SVM) with a linear kernel (*soft u*: 41.69±18.68%, *hard u*: 34.83±14.51%, *p shared*: 32.03±12.54%, and *f shared*: 34.24±16.35% across all time points; Fig. 6b). Regardless of decoding framework, we found that *soft u* consistent spikes are more decodable and potentially more readable downstream.

To ensure that decoding accuracy depended on correspondence between spikes and edges in each sub-FN we permuted the edges of each sub-FN before selecting the subset of action potentials. This allowed us to preserve the density of each sub-FN, as well as the associated spike sparsity (Methods). In all cases decoding performance decreased when selecting spikes associated with permuted edges: performance was 0.57±0.31 and 0.50±0.23 of the real decoding performance of *hard u* and *soft u*, respectively, while the shared sub-FN showed a lesser degradation (0.76±0.33 and 0.66±0.37 for *p shared* and *f shared*, respectively. Fig. 6c).

## Discussion

Numerous studies have demonstrated that pairwise correlations between neurons can be segmented according to the variable that best explains them, including stimulus ^20–22^, local integration ^23,24^ and global or broadcast signals within neocortex^25,26^. This suggested the possibility that pairs of spikes, which correspond to these correlations, would similarly be separable. We segmented spikes according to whether they corresponded to one of four broad classes of pairwise correlations, or sub-FNs. We found that one class of stimulus specific correlation identified sets of spikes, far fewer than the whole, that are particularly informative of visual stimulus. This was the case when measuring information using information theoretic measures and when we evaluated the decodability of the spikes regardless of which class of decoding algorithm we applied. Both as the individual neuron and population levels these informative spikes were not composed of reproducible patterns. Rather, they exhibited comparable levels of trial-to-trial variability to any other set of spikes. Nonetheless, these spikes outperformed all other sets of spikes, including the full set of spikes in every measure tested. Together these finding suggest that reliable pairwise correlations, rather than simply rate modulation or temporal pattern, are the building blocks of the coding scheme employed in layer 2/3 of cortex. A similar coding scheme termed ‘ensemble cofiring’ has been recently proposed in area CA1 in hippocampus^40^.

While we identified pairs of spikes according to their congruence with reliable trial-averaged correlations, which are inaccessible to downstream targets, this coding scheme does not depend *a priori* on this trial averaged knowledge. Rather the suggested single trial coding scheme only requires that neuron pairs with stimulus-specific pairwise correlations between them project to the same downstream target whereas neuron pairs with non-stimulus-specific correlation project to another. Such a connectivity pattern allows dynamics to be disentangled despite overlapping neuron identities and spikes on short timescales. In line with this suggestion, cortical pyramidal neurons have been found to project to multiple targets^41^. Yet, such specific connectivity patterns from layer 2/3 to downstream targets still need to be demonstrated and provide a clear prediction and test^36^ for a pairwise correlation coding scheme.

Two features of the proposed code are especially appealing; first, the single trial coding scheme allows for multiplexing, that is, the population is capable of simultaneously representing multiple variables. The high complexity of the natural world and the fact that sensory and motor related brain activity is broadly distributed^26,42^ argue for multiplexity. Multiplexing has previously been considered for the format of the coding scheme^43–45^, the content of the code (i.e. the feature), such as reported here, or both^46,47^. Notably, format multiplexing, such as rate versus temporal schemes, potentially requires differing readout mechanisms. Our hypothesis calls for a simpler, uniform readout system, based on wiring and correlation rather than cellular properties. While in our work no variables other than the direction of drifting gratings were systematically examined, we postulate that stimulus-nonspecific (shared) correlations might code for other variables, such as internal state. Experiments in which several variables are controlled can thus be used to test our theory empirically.

A second advantage aspect of the coincident-spikes scheme proposed here is that it does not rely on perfectly reproducible patterns of activity across trials and is viable during single trials. This is apparent in soft u consistent spikes containing large information quantity and being accurately decodable despite high trial-to-trial variability of spike counts, temporal patterns and population vectors. Indeed, our findings that trial-to-trial variability in single neurons and population, does not constitute an obstacle for the sensory code. This type of coding scheme, where activity pattern of different structure have similar meaning has been termed a ‘semantic code’^48,49^. In addition to their flexibility and learnability^50^, a semantic coding scheme allows extraction of meaning from sparse firing, as we found here. In turn, sparse coding has been hypothesized to increase the coding capacity of the network and to be metabolically efficient^51,52^.

In summary, here we use functional networks to isolate pairs of spikes that occur during a single trial in a stimulus dependent manner. We find that only a small subset of these pairwise correlations and the corresponding pairs of spikes occur at any given single point in time, resulting in very sparse and variable dynamics during single trials. Nonetheless, these sparse sets of spikes carry more information about the stimulus than all the spikes recorded during the same single trial. Our work highlights the importance of considering the perspective of a downstream reader^36^ when analyzing spike trains, and promotes a spike-centric rather than neuron-centric view of the sensory code.

## Methods

### Data collection and curation

Animals and protocols are described in full in ref 32. We performed a craniotomy over the left primary visual cortex (V1) in 8 Tg(Thy1-GCaMP6s)GP4.12Dkim (Jackson Laboratory) mice (4 male, 4 female). These mice constitutively express GCaMP6s in excitatory pyramidal neurons of layer 2/3. Upon recovery and verification of V1 location, mice were head-fixed but free to run on a linear treadmill while passively viewing drifting gratings in 12 evenly-spaced directions (80% contrast, 0.04 cyc/deg spatial frequency and 2Hz temporal frequency). Stimulus presentation (ON epochs) were 5s in duration and interleaved with 3s of gray screen (OFF epochs). Imaging of the neuronal population was performed with a line-scan^53^ at a wavelength of 910nm (Coherent Chameleon). We inferred spikes from df/f by employing a deconvolution algorithm^54^. All data presented are from 19 datasets unless stated otherwise.

### Functional networks

We summarized the pairwise relationships between neurons in a static functional network (FN), computed using the confluent mutual information of the spikes between every pair of neurons *i,j*. Confluent mutual information (conMI^33^, Fig. 1a) is a non-symmetric value defined as:

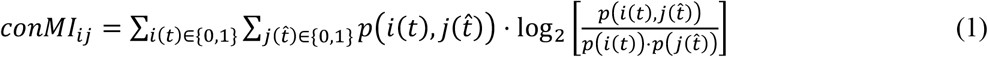

where:

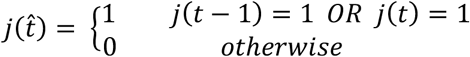

This results in an *N*x*N* adjacency matrix, with neurons being nodes and conMI values as directed and weighted edges between them. Edges between neuronal pairs with negative correlation between their spike trains were then set to zero, and the adjacency matrix was further pruned to contain only the top 50% of edge weights, resulting in a final density of 0.38±0.04. Functional network construction was performed separately for trials of each direction of drifting grating, for a total of 12 FNs per dataset.

### Edge classification into four sub-FNs

For each direction *θ* of drifting gratings, 3 fellow directions were defined: the two adjacent directions that are 30° apart from *θ*, and the opposite direction 180° apart from *θ*, which has the same orientation as *θ*. For example, for *θ*=60°, the fellow directions are 30°, 90° and 240°. We then compared each edge in the FN for *θ* to the values in the three fellow directions according to the following rules:

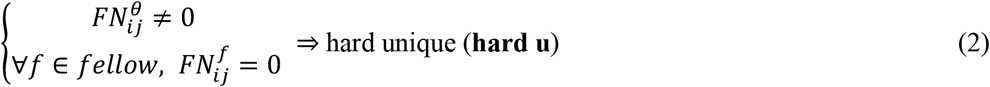

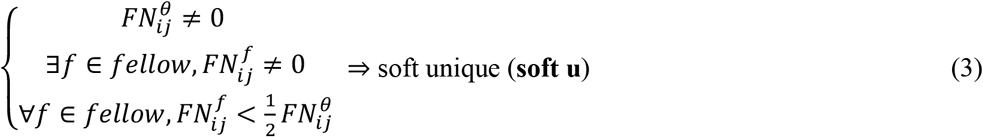

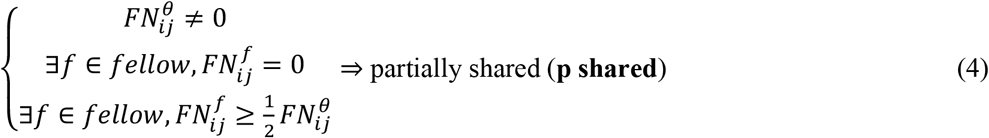

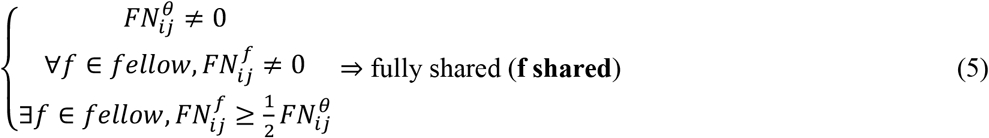

This procedure was carried out for every direction of drifting gratings, resulting in 4 mutually exclusive sub-functional networks (sub-FNs) for each of the 12 directions in each dataset.

### Spike sparsification

A temporal graph (TG) is a single-trial moment-to-moment representation of which pairwise relationships are instantiated. To construct a temporal graph, we first built a binary tensor of potential edges (POT) of size *N*x*N*x*(T-1)* with *T* being the time points in the rasters (*R*). Each time slice summarized the spiking activity at *t-1* and *t*. For each neuron pair *i,j* at each time point sliding along the duration of the spiking activity:

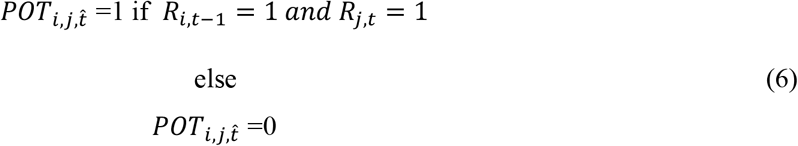

POT is called a tensor of potential edges since it may be the case that *i* spiking at *t-1* has contributed to the spiking activity of *j* at *t*. POT is thus a combinatorial representation of the rasters and its density depends on the firing rate in the population.

Each time slice was then intersected with an FN, or sub-FN, as indicated in the results, to give a temporal graph (TG):

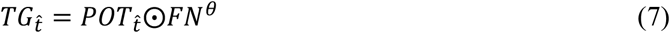

A TG is thus the same size as the corresponding potential edges tensor, but is sparser and contains weighted edges.

Then, for every neuron pair *i,j* and every time slice, we defined the sparsified rasters (SR) to be:

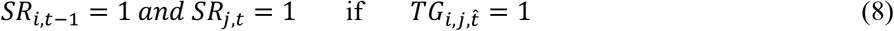

Spikes in SR are a subset of the spikes in the original raster R that can be explained as a manifestation of the FN or the sub-FN that was used in the calculation of the TG (Fig. 3a).

### Trial-to-trial variability measures

Single cell rate variability (Fig. 4), was the coefficient of variation, CV as *σ/μ* with *σ* and *μ* being the standard deviation and mean, respectively, of the neuron spike-count across trials with the same drifting gratings direction. For temporal precision of single cells, for each neuron we calculated the Victor-Purpura (VP) distance^6^ between pairs of trials of the same direction. Since VP is sensitive to the number of spikes (i.e. sparsity), we divided by the mean spike count for the trials in the pair. We chose *q*=1 for the cost. Population-level variability was measured by the L2 (Euclidean) norm between each pair of population vectors for trials of the same drifting grating direction. We formed population vectors by counting the spikes for each neuron across the duration of the trial. The L2 metric was normalized by the mean of the total spikes in the trials in the pair to adjust for sparsity differences.

### Information quantification

The mutual information (in bits) between the neural response r and the stimulus directions was calculated as^55^:

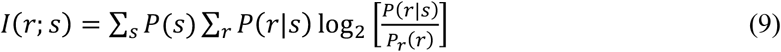

The neural responses *r* were binarized vectors of spikes for 5 neurons binned across 10 imaging frames (Fig. 5a), and vectors of spikes for 5 neurons across 3 imaging frames vertically concatenated into 15-by-1 vectors (Fig. 5b). For all sub-FN consistent spikes in each dataset, we selected the 5 neurons with the highest (average, in Fig. 5c) firing rate across the first 100 imaging frames of all trials in the original spikes. The above sum is then computed across all unique values of *r* for each stimulus direction *s*, using *P(s) =1/12* for all *s*, and we scaled raw information value by the average number of spikes in a vector to obtain bits per spike. In Fig. 5d we have used the same 5 neurons but took *r* to be spikes that are *not* consistent with edges in the sub-FN examined.

### Feedforward pattern recognition neural network decoder

We employed a multiclass decoder with N input units for N neurons, and 12 output units, one for each direction of drifting gratings. The input and output layer are connected by all-to-all feedforward connectivity with random initial weights. As inputs, we binned spikes in 10 consecutive imaging frames to a population vector of size N. 90%-10% of the trials were used as training and test sets, respectively. The weights were trained by conjugate gradient with Matlab’s machine learning toolbox. We trained the weights for the original spikes and the sub-FN consistent spikes separately.

### Support vector machine (SVM) decoder

To confirm the results of the feedforward neural network decoder, we used the same inputs, i.e. population vectors of spikes binned at 10 frames from 90% of the trials, to train a Support Vector Machine (SVM) decoder with a linear kernel. Training was performed with Matlab’s fitcecoc.m function. We tested on the remaining 10% of trials.

### Controls

In order to test for the possibility that any spiking activity that is organized in pairs of consecutive spikes accounts for the decoding performance, we designed a stringent control that preserves the density of the sub-FNs as well as the resulting pairwise structure of spikes. We randomly permuted the edges in each sub-FN before intersecting with each trial. We then binned the spikes in each 10 frames as with the real spikes, and passed them through the SVM decoder as described above.

### Statistical analysis and code

Means and standard deviations across datasets or neurons are reported throughout the paper as M±SD unless stated otherwise. In the case of analysis with multiple time points (for example, Fig 6a,b and Fig 7a-c) analysis of variance (ANOVA) was performed for every time point separately. p values were bonferroni corrected in cases of multiple comparisons. All analysis was done in Matlab 2018 or later (Mathworks) and Python 3.7.

## References

1. Niell, C. M. & Stryker, M. P. Modulation of visual responses by behavioral state in mouse visual cortex. Neuron 65, 472–479 (2010).

2. Stringer, C. et al. Spontaneous behaviors drive multidimensional, brainwide activity. Science 364, (2019).

3. Hubel, D. H. & Wiesel, T. N. Receptive fields of single neurones in the cat’s striate cortex. J. Physiol. 148, 574–591 (1959).

4. Hubel, D. H. & Wiesel, T. N. Receptive fields and functional architecture of monkey striate cortex. J. Physiol. 195, 215–243 (1968).

5. Phillips, D. P. & Irvine, D. R. Responses of single neurons in physiologically defined primary auditory cortex (AI) of the cat: frequency tuning and responses to intensity. J. Neurophysiol. 45, 48–58 (1981).

6. Victor, J. D. & Purpura, K. P. Nature and precision of temporal coding in visual cortex: a metric-space analysis. J. Neurophysiol. 76, 1310–1326 (1996).

7. Chi, Z. & Margoliash, D. Temporal Precision and Temporal Drift in Brain and Behavior of Zebra Finch Song. Neuron 32, 899–910 (2001).

8. Rullen, R. V. & Thorpe, S. J. Rate Coding Versus Temporal Order Coding: What the Retinal Ganglion Cells Tell the Visual Cortex. Neural Comput. 13, 1255–1283 (2001).

9. Ventura, V. Trial-to-Trial Variability and Its Effect on Time-Varying Dependency Between Two Neurons. J. Neurophysiol. 94, 2928–2939 (2005).

10. Tolhurst, D. J., Movshon, J. A. & Dean, A. F. The statistical reliability of signals in single neurons in cat and monkey visual cortex. Vision Res. 23, 775–785 (1983).

11. Gur, M., Beylin, A. & Snodderly, D. M. Response variability of neurons in primary visual cortex (V1) of alert monkeys. J. Neurosci. Off. J. Soc. Neurosci. 17, 2914–2920 (1997).

12. Deweese, M. R. & Zador, A. M. Shared and Private Variability in the Auditory Cortex. J. Neurophysiol. 92, 1840–1855 (2004).

13. Shadlen, M. N. & Newsome, W. T. The variable discharge of cortical neurons: implications for connectivity, computation, and information coding. J. Neurosci. Off. J. Soc. Neurosci. 18, 3870–3896 (1998).

14. Osborne, L. C., Palmer, S. E., Lisberger, S. G. & Bialek, W. The Neural Basis for Combinatorial Coding in a Cortical Population Response. J. Neurosci. 28, 13522–13531 (2008).

15. Luczak, A., McNaughton, B. L. & Harris, K. D. Packet-based communication in the cortex. Nat. Rev. Neurosci. 16, 745–755 (2015).

16. Churchland, M. M. et al. Neural population dynamics during reaching. Nature (2012) doi:10.1038/nature11129.

17. Averbeck, B. B., Latham, P. E. & Pouget, A. Neural correlations, population coding and computation. Nat. Rev. Neurosci. 7, 358–366 (2006).

18. Cohen, M. R. & Kohn, A. Measuring and interpreting neuronal correlations. Nat. Neurosci. 14, 811–819 (2011).

19. Salinas, E. & Sejnowski, T. J. Correlated neuronal activity and the flow of neural information. Nat. Rev. Neurosci. 2, 539–550 (2001).

20. Levy, M., Sporns, O. & MacLean, J. N. Network Analysis of Murine Cortical Dynamics Implicates Untuned Neurons in Visual Stimulus Coding. Cell Rep. 31, 107483 (2020).

21. Stevenson, I. H. et al. Functional Connectivity and Tuning Curves in Populations of Simultaneously Recorded Neurons. PLOS Comput. Biol. 8, e1002775 (2012).

22. Ponce-Alvarez, A., Thiele, A., Albright, T. D., Stoner, G. R. & Deco, G. Stimulus-dependent variability and noise correlations in cortical MT neurons. Proc. Natl. Acad. Sci. 110, 13162–13167 (2013).

23. Chambers, B. & MacLean, J. N. Higher-Order Synaptic Interactions Coordinate Dynamics in Recurrent Networks. PLOS Comput. Biol. 12, e1005078 (2016).

24. Salinas, E. & Sejnowski, T. J. Impact of correlated synaptic input on output firing rate and variability in simple neuronal models. J. Neurosci. Off. J. Soc. Neurosci. 20, 6193–6209 (2000).

25. Ecker, A. S., Berens, P., Tolias, A. S. & Bethge, M. The Effect of Noise Correlations in Populations of Diversely Tuned Neurons. J. Neurosci. 31, 14272–14283 (2011).

26. Musall, S., Kaufman, M. T., Juavinett, A. L., Gluf, S. & Churchland, A. K. Single-trial neural dynamics are dominated by richly varied movements. Nat. Neurosci. 22, 1677–1686 (2019).

27. Ganmor, E., Segev, R. & Schneidman, E. Sparse low-order interaction network underlies a highly correlated and learnable neural population code. Proc. Natl. Acad. Sci. 108, 9679–9684 (2011).

28. Pillow, J. W. et al. Spatio-temporal correlations and visual signalling in a complete neuronal population. Nature 454, 995–999 (2008).

29. Kotekal, S. & MacLean, J. N. Recurrent interactions can explain the variance in single trial responses. PLOS Comput. Biol. 16, e1007591 (2020).

30. Harris, K. D. Neural signatures of cell assembly organization. Nat. Rev. Neurosci. 6, 399–407 (2005).

31. Köster, U., Sohl-Dickstein, J., Gray, C. M. & Olshausen, B. A. Modeling Higher-Order Correlations within Cortical Microcolumns. PLoS Comput. Biol. 10, e1003684 (2014).

32. Dechery, J. B. & MacLean, J. N. Functional triplet motifs underlie accurate predictions of single-trial responses in populations of tuned and untuned V1 neurons. PLOS Comput. Biol. 14, e1006153 (2018).

33. Chambers, B., Levy, M., Dechery, J. B. & MacLean, J. N. Ensemble stacking mitigates biases in inference of synaptic connectivity. Netw. Neurosci. 1–49 (2017).

34. Cossell, L. et al. Functional organization of excitatory synaptic strength in primary visual cortex. Nature 518, 399–403 (2015).

35. Ko, H. et al. The emergence of functional microcircuits in visual cortex. Nature 496, 96–100 (2013).

36. Buzsáki, G. Neural syntax: cell assemblies, synapsembles and readers. Neuron 68, 362–385 (2010).

37. Shannon, C. E. A mathematical theory of communication. Bell Syst. Tech. J. 27, 379–423 (1948).

38. Strong, S. P., Koberle, R., de Ruyter van Steveninck, R. R. & Bialek, W. Entropy and Information in Neural Spike Trains. Phys. Rev. Lett. 80, 197–200 (1998).

39. Dan, Y., Alonso, J.-M., Usrey, W. M. & Reid, R. C. Coding of visual information by precisely correlated spikes in the lateral geniculate nucleus. Nat. Neurosci. 1, 501–507 (1998).

40. Levy, E. R. J., Park, E. H., Redman, W. T. & Fenton, A. A. A neuronal code for space in hippocampal coactivity dynamics independent of place fields. bioRxiv (2021). doi:10.1101/2021.07.26.453856.

41. Kasthuri, N. et al. Saturated Reconstruction of a Volume of Neocortex. Cell 162, 648–661 (2015).

42. Steinmetz, N. A., Zatka-Haas, P., Carandini, M. & Harris, K. D. Distributed coding of choice, action and engagement across the mouse brain. Nature 576, 266–273 (2019).

43. Naud, R. & Sprekeler, H. Sparse bursts optimize information transmission in a multiplexed neural code. Comput. Biol. 10.

44. Kayser, C., Montemurro, M. A., Logothetis, N. K. & Panzeri, S. Spike-phase coding boosts and stabilizes information carried by spatial and temporal spike patterns. Neuron 61, 597–608 (2009).

45. Ainsworth, M. et al. Rates and Rhythms: A Synergistic View of Frequency and Temporal Coding in Neuronal Networks. Neuron 75, 572–583 (2012).

46. Lankarany, M., Al-Basha, D., Ratté, S. & Prescott, S. A. Differentially synchronized spiking enables multiplexed neural coding. Proc. Natl. Acad. Sci. 116, 10097–10102 (2019).

47. Insanally, M. N. et al. Spike-timing-dependent ensemble encoding by non-classically responsive cortical neurons. Elife 8, e42409 (2019).

48. Ganmor, E., Segev, R. & Schneidman, E. A thesaurus for a neural population code. Elife 4, e06134 (2015).

49. Granot-Atedgi, E., Tkačik, G., Segev, R. & Schneidman, E. Stimulus-dependent Maximum Entropy Models of Neural Population Codes. PLOS Comput. Biol. 9, e1002922 (2013).

50. Berry, M. J. I. & Tkačik, G. Clustering of Neural Activity: A Design Principle for Population Codes. Front. Comput. Neurosci. 14, (2020).

51. Vinje, W. E. & Gallant, J. L. Sparse Coding and Decorrelation in Primary Visual Cortex During Natural Vision. Science (2000). doi:10.1126/science.287.5456.1273.

52. Olshausen, B. A. & Field, D. J. Sparse coding with an overcomplete basis set: A strategy employed by V1? Vision Res. 37, 3311–3325 (1997).

53. Sadovsky, A. J. et al. Heuristically optimal path scanning for high-speed multiphoton circuit imaging. J. Neurophysiol. 106, 1591–1598 (2011).

54. Friedrich, J., Zhou, P. & Paninski, L. Fast online deconvolution of calcium imaging data. PLoS Comput. Biol. 13, e1005423 (2017).

55. Brenner, N., Strong, S. P., Koberle, R., Bialek, W. & Steveninck, R. R. de R. van. Synergy in a Neural Code. Neural Comput. 12, 1531–1552 (2000).

